# RNA-Dependent Amplification of Mammalian mRNA Encoding Extracellular Matrix Components: Identification of Chimeric RNA Intermediates for α1, β1, and γ1 Chains of Laminin

**DOI:** 10.1101/376293

**Authors:** Vladimir Volloch, Sophia Rits, Bjorn Olsen

## Abstract

The aim of the present study was to test for the occurrence of key elements predicted by the previously postulated mammalian RNA-dependent mRNA amplification model in a tissue producing massive amounts of extracellular matrix proteins. At the core of RNA-dependent mRNA amplification, until now only described in one mammalian system, is the self-priming of an antisense strand and extension of its 3’ terminus into a sense-oriented RNA containing the protein-coding information of a conventional mRNA. The resulting product constitutes a new type of biomolecule. It is chimeric in that it contains covalently connected antisense and sense sequences in a hairpin configuration. Cleavage of this chimeric intermediate in the loop region of a hairpin structure releases mRNA which contains an antisense segment in its 5’UTR; depending on the position of self-priming, the chimeric end product may encode the entire protein or its C-terminal fragment. The occurrence of such composite chimeric molecules is unique for this type of mRNA amplification and represents a conclusive “identifier” of this process. We report here the detection, by next generation sequencing, of such chimeric junction sequences for mRNAs molecules encoding αl, β1, and γ1 chains of laminin in cells of the extracellular matrix-generating Engelbreth-Holm-Swarm (EHS) mouse tumor, best known for producing extraordinarily large amounts of “Matrigel”.

## INTRODUCTION

Previously, we described evidence in support of RNA-dependent amplification of mammalian mRNA (1, 2). This amplification, of alpha- and beta-globin-encoding mRNA, occurred during terminal erythroid differentiation and, therefore, in cells destined to die in a few days. For this reason, it could be argued that the amplification mechanism might be in restricted use and not relevant to normal tissue formation and functioning. Therefore, assessing the scope of the contribution of this cellular mechanism to development and homeostasis of different tissues is important in establishing its physiological significance. Among potential cell types that may utilize mRNA amplification by the mechanism described above are cells producing extracellular matrix proteins. At several stages during the development, growth and repair of connective tissues, matrix proteins are rapidly produced and, because they are secreted, the production of extraordinary amounts is not limited by cellular confines. A mechanism, described in detail in the Discussion section below, whereby every mRNA molecule may serve as a potential template for production of additional mRNA, may facilitate increased production of specific secreted proteins at certain times. With this in mind, the present study utilized cells from the Engelbreth-Holm-Swarm mouse sarcoma tumor that generate, in extraordinarily large quantities, basement membrane components that are collected and marketed as “Matrigel” (3–10).

The transfer of information in postulated mRNA amplification is mediated by RNA-dependent RNA polymerase (RdRp) via synthesis of antisense RNA and is diagrammatically illustrated in Figure 1.

**Figure 4.**
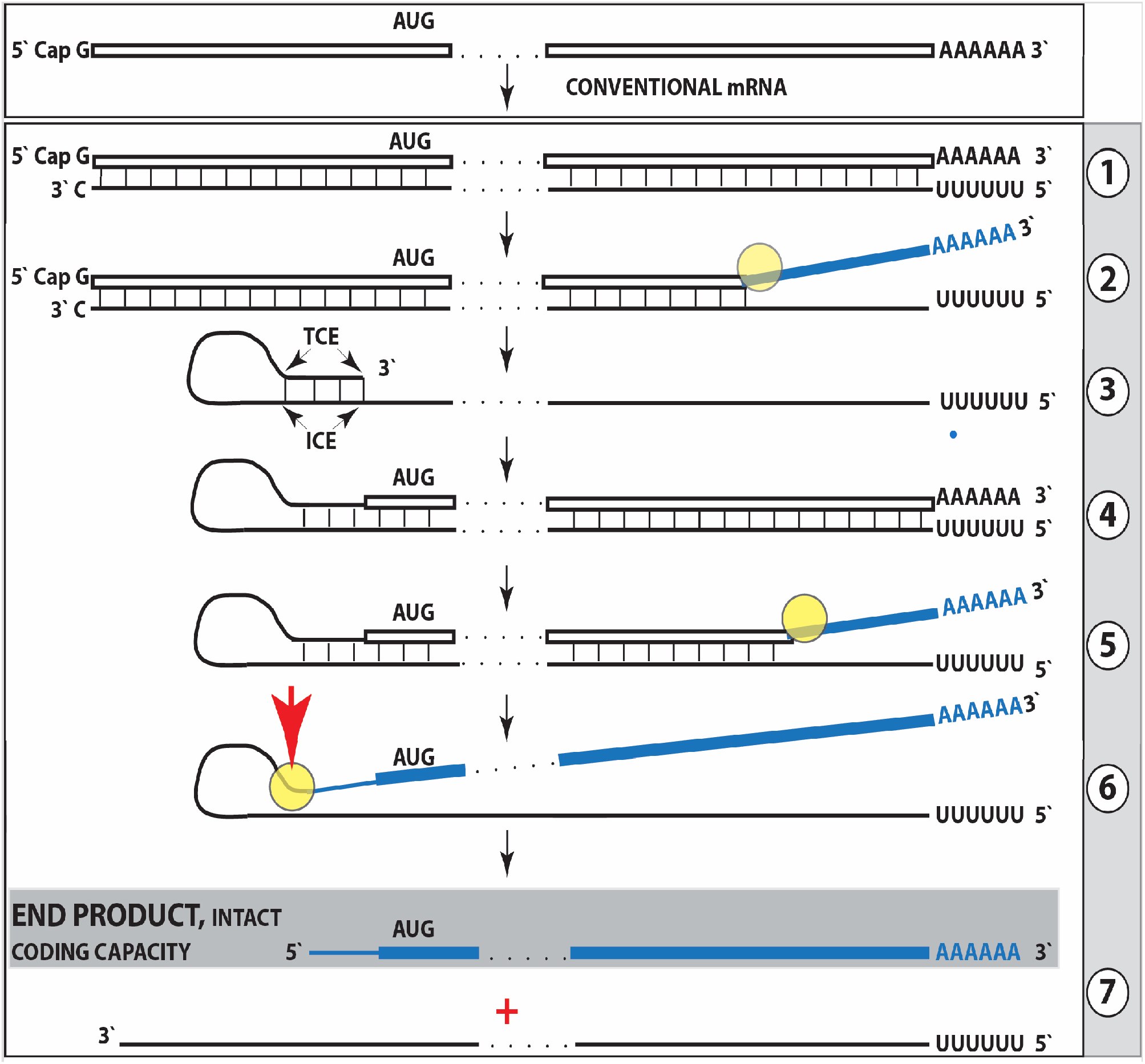
Projected steps in RdRp- and antisense-mediated amplification of mRNA. <u>Top panel</u>: Conventional, genome-transcribed mRNA molecule. Bottom panel: postulated stages of antisense-mediated mRNA amplification. Boxed line – sense strand RNA. Single line – antisense strand RNA. “AUG” – functional translation initiation codon. “TCE”– 3’-terminal complementary element; “ICE”– internal complementary element, both on the antisense strand. Yellow circle – helicase/modifying activity complex. Blue lines (both single and boxed) – RNA strand modified and separated from its complement by a helicase complex. Red arrowhead – position of cleavage of the chimeric intermediate. Step 1: synthesis of antisense strand; step 2: strand separation; step 3: folding of antisense strand into self-priming configuration; step 4: extension of self-primed antisense RNA; step 5: strand separation; step 6: cleavage of the chimeric intermediate; stage 7: end-products of amplification. For more details please see the Discussion section.

Briefly, synthesis of antisense strand would start at the 3’-poly(A) of an mRNA molecule, initiated, possibly, with a uridylated protein as suggested by observations in the poliovirus system (25). After separation from its template, the 3’-terminal region of the antisense strand self-primes its extension into sense-oriented RNA strand. The self-priming of the antisense molecule requires the presence of sequences, described in detail in the Discussion section below, that enable the amplification process and define its specificity. The completion of the extension results in a new type of biomolecule: a chimeric RNA molecule containing both antisense and sense sequences in a hairpin configuration. This molecule is the immediate precursor of amplified mRNA which may be released by the cleavage at the loop of a hairpin structure and separation of sense and antisense strands. The occurrence of composite chimeric molecules is unique for this process. Therefore, the presence of chimeric antisense/sense RNA junction sequences serves as a conclusive identifier of the occurrence of critical initial steps in mRNA amplification. We report below the detection, by next generation sequencing, of such definitive identifiers, namely the chimeric sense/antisense junction sequences corresponding to Steps 4 and 5 of Figure 1, for mRNAs encoding α1, β1, and γ1 chains of laminin in “Matrigel”-producing cells.

## RESULTS

The occurrence of chimeric junctions containing both antisense and sense sequences in mRNAs encoding ECM components produced by EHS tumor cells was probed by next generation sequencing (NGS). For two reasons, experiments described below were carried out with cytoplasmic RNA. First, although it would be more convenient to isolate and work with total cellular RNA, we could, as described in the Methods section, loose some molecules of interest in the process. Second, there is good reason to believe that molecules of interest, if they are indeed intermediates in an mRNA amplification process, are located in the cytoplasm. This is because the enzyme involved, mammalian RNA-dependent RNA polymerase has been found in the cytoplasm but not in the nucleus (20). Cytoplasmic RNA was isolated from tumor cells harvested ten days after tumor implantation, and, following depletion of ribosomal RNA, used to generate sequencing libraries. Libraries were sequenced on an Illumina Next500 instrument. Sequencing data were converted into blast databases and analyzed by blasting with chimeric reference sequences. Sequences of interest were extracted from raw data and analyzed. Contiguous chimeric sequences containing antisense and sense components with the junction occurring within 5’ untranslated regions of mRNA species encoding α1, β1, and γ1 chains of laminin were detected. Complete fragments included the following sequences (lowercase: antisense orientation; uppercase: sense orientation) that served as a basis for Figures 2-4 shown below:

> Laminin α1, fragment1 (Figure 2, top panel):
>
> cgcccgtgctcgctcacaggcactgcgcgagtcCTTCCCCAGGAGCGCAGGGAGCGGCGGCGACAACATGCGCGGCAGCGGCACGGGAGCCGCGCTCCTGGTGCTCCTGG.

> Laminin α1, fragment 2 (Figure 2, bottom panel):
>
> ggtgacccagagcaccgaggccaggagcaccaggagcgcggctcccgtgccgctgccgcgcatgttgtcgccgccgctccctgcgctcctggggaaggtctcgcccgtgctcgctcacaggcactgcgcgagtgtgctcttccCCAGGAGCGCAGGGAGCGGC.

> Laminin β1, fragment 1 (Figure 3, top panel):
>
> ggcccttccatttcctgccgctcccacggaagcgggggctgggcccaggaagggggtcaagtcgcttaactactttgttctcctcacccggctgggcgagcgctcaacccgctcctggcagcccaccggGTGAGGAGAACAAAGTAGTTAAGCGAC.

> Laminin β1, fragment 2 (Figure 3, bottom panel):
>
> tgggcccaggaagggggtcaagtcgcttaactactttgttctcctcacccggctgggcgagcgctcaacccgctcctggcagcccaccggccccCGCTTCCGTGGGAGCGGCAGGAAATGGAAGGGCCCC.

> Laminin γ1, fragment 1 (Figure 4, top panel):
>
> cggtgcgcagcctgcactgtgcgccgGGAGGTAGCGGAGGCAGCGCGATCTTGGCTCGGACGCCCACCCATCGGCTCTGCGTCCGGCTCTCGGCCTCCAGCCCGGTCCACAGCCCGGCCTCGGCCCGCAGCGGAGGATCGGCCTCGGGATACGCCGCTAG.

> Laminin γ1, fragment 2 (Figure 4, bottom panel):
>
> ccgacttccgagcgcgcactcgagagcgcgcTCGGAAGTCGGGGGTCGGCGCGCAGTGCAGGCTGCGCA
>
> CCGGGAGGTAGCGGAGGCAGCG.

**Figure 2.**
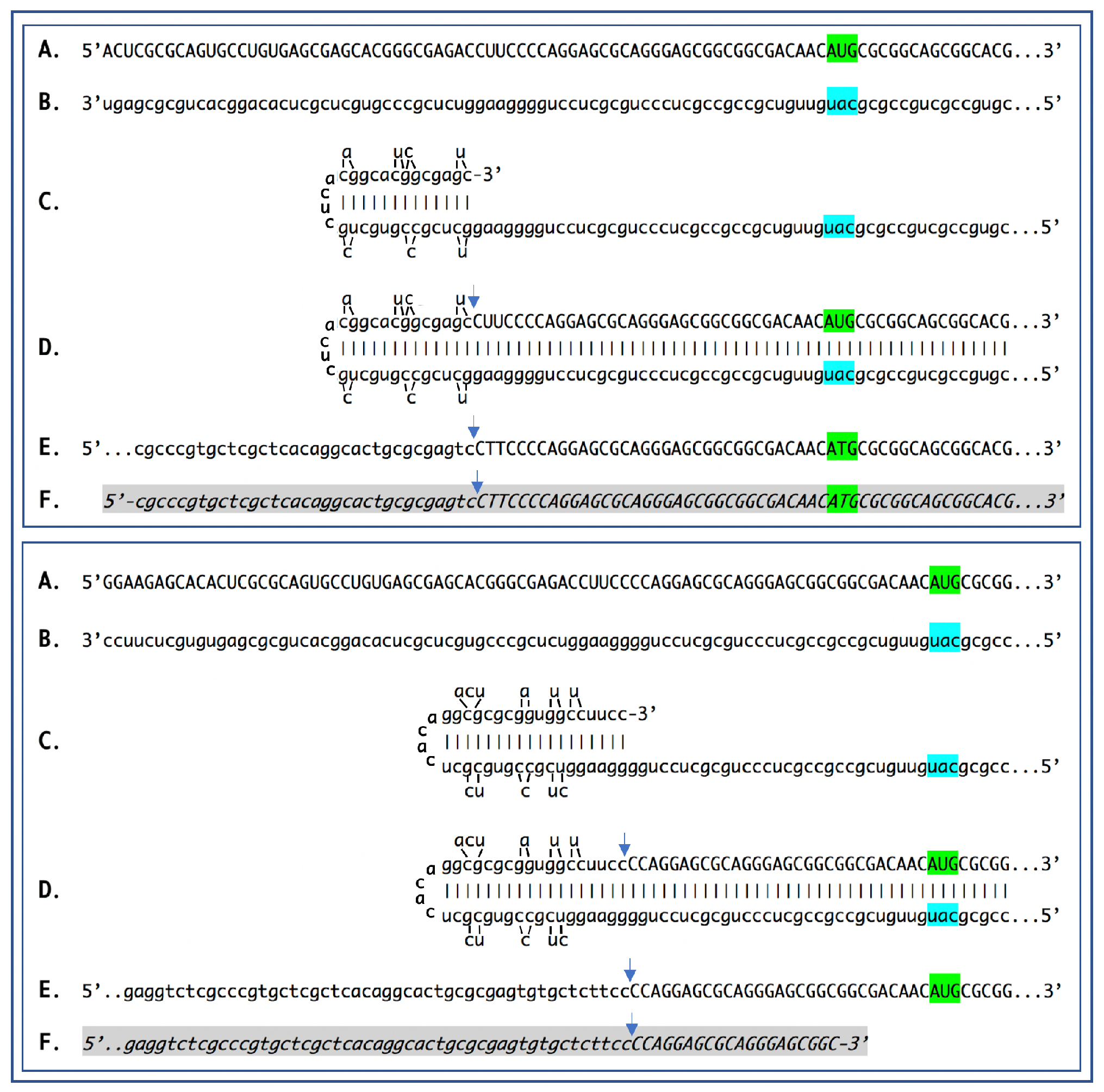
Sequence of a chimeric junction containing antisense and sense segments of laminin α1 mRNA and the projected pathway of its generation. Uppercase letters – nucleotide sequence of the sense strand; lowercase letters – nucleotide sequence of the antisense strand. Highlighted in green – “AUG” translation initiation codon on the sense strand; highlighted in blue – “uac” complement of translation initiation codon on the antisense strand. In italics and highlighted in grey – detected chimeric fragments. Blue arrows: position of antisense/sense junction. The top and bottom panels depict amplification of mRNA molecules originated from different TSSs; note that self-priming positions and, consequently, chimeric junction sequences in two panels are different. **A**: 5’ terminus of laminin α1 mRNA. **B**: antisense complement of the 5’ terminus of α1 laminin mRNA. **C**: folding of the antisense strand into self-priming configuration. **D**: extension of selfprimed antisense strand into sense-oriented sequence. **E**: projected chimeric junction sequence. **F**: detected chimeric junction sequence. Complete sequences of the detected chimeric fragments are provided in the main text above. Note that the priming occurs within the segment of antisense strand corresponding to the 5’UTR of mRNA, thus preserving the coding capacity of amplified mRNA.

**Figure 3.**
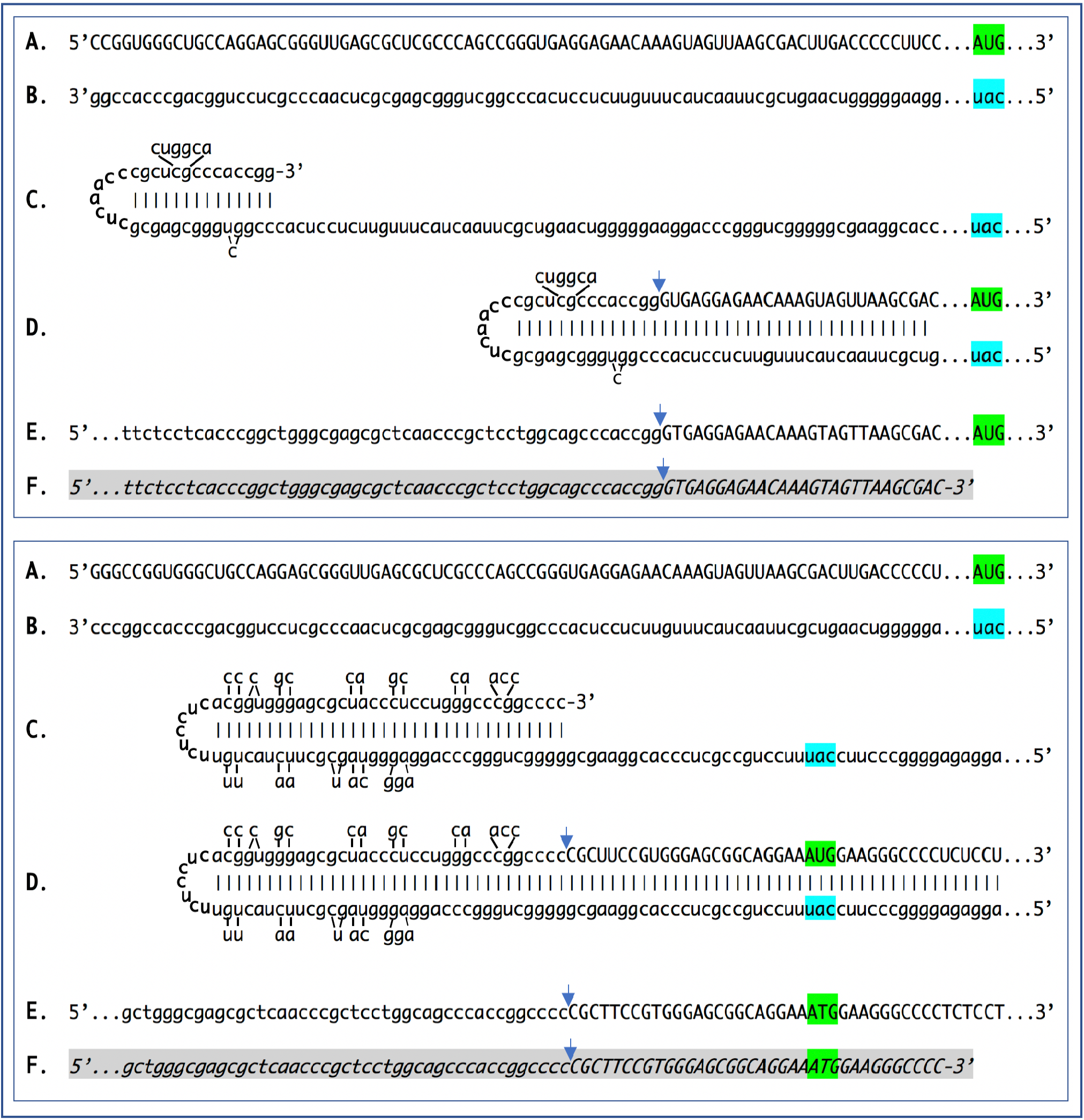
Sequence of a chimeric junction containing antisense and sense segments of laminin β1 mRNA and the projected pathway of its generation. Uppercase letters – nucleotide sequence of the sense strand; lowercase letters – nucleotide sequence of the antisense strand. Highlighted in green – “AUG” translation initiation codon on the sense strand; highlighted in blue – “uac” complement of translation initiation codon on the antisense strand. In italics and highlighted in grey – detected chimeric fragments. Blue arrows: position of sense/antisense junction. The top and bottom panels depict amplification of mRNA molecules originated from different TSSs; note that self-priming positions and, consequently, chimeric junction sequences shown in two panels are different. **A**: 5’ terminus of laminin β1 mRNA. **B**: antisense complement of the 5’ terminus of a1 laminin mRNA. **C**: folding of the antisense strand into self-priming configuration. **D**: extension of self-primed antisense strand into sense-oriented sequence. **E**: projected chimeric junction sequence. **F**: detected chimeric junction sequence. Complete sequences of the detected chimeric fragments are provided in the main text above. Note that the priming occurs within the segment of antisense strand corresponding to the 5’UTR of mRNA, thus preserving the coding capacity of amplified mRNA.

**Figure 4.**
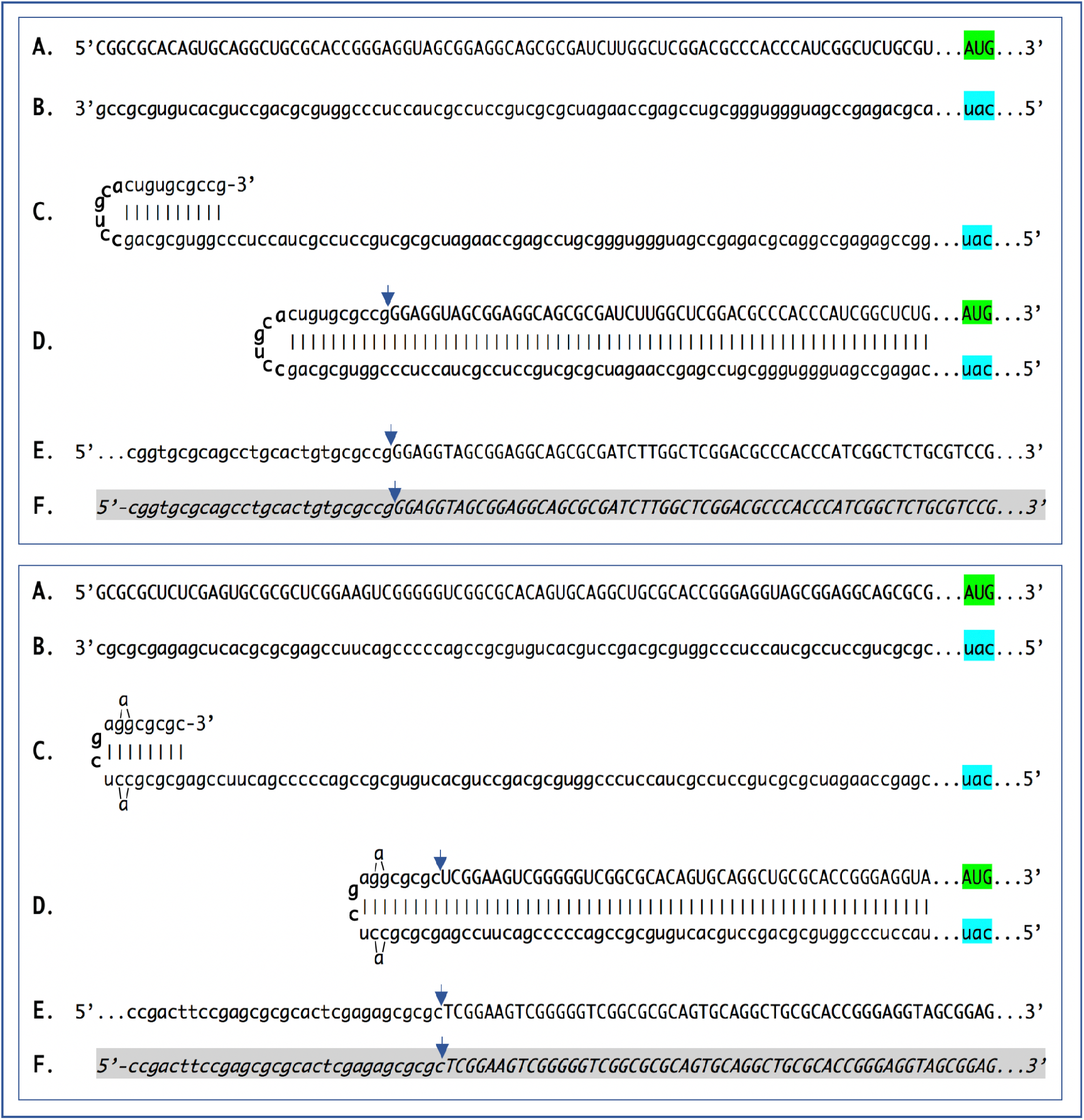
Sequence of a chimeric junction containing antisense and sense segments of laminin γ1 mRNA and the projected pathway of its generation. Uppercase letters – nucleotide sequence of the sense strand; lowercase letters – nucleotide sequence of the antisense strand. Highlighted in green – “AUG” translation initiation codon on the sense strand; highlighted in blue – “uac” complement of translation initiation codon on the antisense strand. In italics and highlighted in grey – detected chimeric fragments. Blue arrows: position of sense/antisense junction. The top and bottom panels depict amplification of mRNA molecules originated from different TSSs; note that self-priming positions and, consequently, chimeric junction sequences shown in two panels are different. **A**: 5’ terminus of laminin γ1 mRNA. **B**: antisense complement of the 5’ terminus of a1 laminin mRNA. **C**: folding of the antisense strand into self-priming configuration. **D**: extension of self-primed antisense strand into sense-oriented sequence. **E**: projected chimeric junction sequence. **F**: detected chimeric junction sequence. Complete sequences of the detected chimeric fragments are provided in the main text above. Note that the priming occurs within the segment of antisense strand corresponding to the 5’UTR of conventional mRNA, thus preserving the coding capacity of amplified mRNA.

The relevant portions of the above sequences are shown in Figures 2-4 together with the projected pathways of their generation detailed in the Figure legends. The sequences are unique in that they are not encoded in the genome contiguously and could be generated only through the extension of the 3’ terminus of a self-primed antisense RNA with the extension start site (ESS), and consequently the chimeric junction sequence, defined precisely and uniquely by the position of a self-priming assembly. Accordingly, chimeric junction sequences serve as identifiers of critical initial steps in the RNA-dependent mRNA amplification process.

The feasibility of the amplification process is largely decided by the sequence and structure of the 3’ termini of the antisense strands, which, in turn, is determined by the position of the transcription start site (TSS) of the gene-encoded mRNA. For all three chains of laminin 111, the chimeric RNA sequences detected originated from mRNAs transcribed from multiple TSS positions. This is not surprising considering that the laminin genes and, in fact, the majority of genes encoding extracellular matrix proteins, lack the “TATA” regulatory element that rigorously defines the position of a TSS. Thus, in the top panel of Figure 2, transcription of the laminin a1 mRNA starts 70 nucleotides upstream from the AUG translation initiation codon. The 3’-terminal region of the corresponding antisense strand is then folded into a self-priming configuration and extended into the sense-oriented component of a chimeric molecule. In the bottom panel of Figure 2, the TSS is shifted ten nucleotides upstream. The antisense molecule forms a different self-priming structure, whose extension from a new ESS yields a distinct chimeric sequence, but the end-result, as defined by the coding content of the amplified sense component, remains unchanged. Indeed, as reasoned in the Discussion section below, the translational outcome of mRNA amplification is defined by the position of self-priming, and because in both cases above, as well as in the cases described below, self-priming occurs within the segment of the antisense strand corresponding to the 5’UTR of the sense strand, the amplified mRNA retains the intact coding capacity of the conventional mRNA.

A similar phenomenon is seen with mRNA for the β1 chain of laminin. Transcription from the TSS position 118 nucleotides upstream from the AUG translation initiation codon produces mRNA whose antisense counterpart is capable of self-priming its extension into a sense strand, thus producing a chimeric molecule (Figure 3, top panel). A shift of the TSS position just three nucleotides upstream results in a radically different self-priming assembly of the antisense strand (Figure 3, bottom panel) which is, nevertheless, extended from a new ESS into a sense strand producing a different chimeric molecule but with the unchanged coding capacity. Similarly, initiation of transcription of mRNA encoding the γ1 chain of laminin from TSS positions 210 nucleotides upstream from the AUG translation initiation codon (Figure 4, top panel) or 246 nucleotides upstream from the translation start site (Figure 4, bottom panel), lead to antisense molecules capable of self-priming their 3’-extension into sense-oriented molecules. The resulting composite antisense/sense molecules have different chimeric junction sequences but retain the protein-coding content identical to that of the conventional mRNA.

## DISCUSSION

The essence of the present study is the detection of a new type of biomolecule. Its structure is evident: a chimeric RNA molecule containing 3’-terminal segment of the antisense strand covalently connected, at a precisely determined position, to the sense strand segment of the same mRNA species. Whereas numerous mechanistic aspects of the overall cellular mechanism involving this type of biomolecule remain to be elucidated and are the subject of future investigations, the mechanism of generation of the chimeric sequences observed in the present study is very clear: the extension of the antisense strand self-primed by its 3’-terminal segment. The position of the junction between the antisense and the sense components of a chimeric molecule is uniquely defined by the folding of the antisense into self-priming configuration. As shown in the present study, this, in turn, is specified by the location of transcription start site(s) of a conventional genome-transcribed mRNA molecule. Moreover, multiple TSSs may lead to multiple chimeric RNAs with junctions at different positions for a single mRNA species. The chimeric fragments described above were identified by next generation sequencing. This procedure not only provides the nucleotide sequence of a single molecule but contains safeguards against detection of chimeric fragments that could be possibly generated during sequencing library preparation. As described below, even if such fragments were generated during the procedure, they could not be sequenced. Thus, unlike any other approach, the procedure used in the present study assures cellular origin of the detected chimeric sequences.

Previously, it was shown that during reverse transcription, provided the antisense strand (cDNA) contains the required complementary elements, self-priming can occur and a fragment containing both sense and antisense components may be generated (14). For several reasons, it can be stated with a high degree of certainty that the chimeric laminin RNA fragments in the present study did not arise during the sequencing library preparation but were present in the initial cytoplasmic RNA preparations. First, any possible synthesis of the second cDNA strand during the stage of first strand synthesis was suppressed by the addition of the NEB strand specificity reagent, as described in the Methods section. Second, even if a hairpin structure were created during the reverse transcription stage, it would represent a dead end in the library preparation process because no adaptor could be ligated. Thirdly, the structure of many of the chimeric fragments detected in the present study decisively rules out the possibility of their generation during the reverse transcription stage of library preparation. This is because the length of the antisense component in these fragments is significantly smaller than that of the sense-oriented segment. Therefore, the former could not serve as a template for the latter during the library preparation process. The only other stage for potential artificial generation of a chimeric fragment, either by intramolecular or intermolecular processes, is the PCR enrichment of a sequencing library. For such an artifact to occur, the 3’-terminal portion of the antisense strand should anneal to the internal portion of the same or another antisense strand and be extended into a sense strand. In sequencing library preparation, the PCR enrichment step follows the adaptor ligation step. If antisense strand contains an adaptor, its 3’-terminal complementary element cannot be extended, even if it anneals either intramolecularly or intermolecularly. If antisense strand doesn’t contain an adaptor, it potentially can anneal and be extended but cannot be sequenced. In this respect, it should also be pointed out that our searches did not detect chimeric fragments for the major housekeeping RNAs. Most importantly as a control, no chimeric fragments were detected for the RNA encoding laminin chains using RNA preparations from an early stage EHS tumor (six days after implantation, as opposed to ten days), where laminin mRNAs are already expressed at high levels but the amplification process, apparently, not yet activated.

The results of the present study show the occurrence of chimeric molecules containing covalently bound sense and antisense strands of RNA encoding all three chains of laminin in a tissue that produces extraordinarily high quantities of this protein. While limited to the demonstration of a new type of biomolecules, these results are indisputable. Likewise is their origin: these molecules are clearly produced by the extension of the 3’terminus of antisense RNA self-primed in a sequence-specific manner. These results can be interpreted within the framework of the mRNA amplification model developed and postulated in more detailed previous studies (1, 2, 11–18) of terminally differentiating cells where the putative mRNA end product of the amplification process accumulates to extraordinary high levels, constituting nearly a tenth of total ribosomal RNA (2). The findings of the present study are fully consistent with the postulated mechanism of mRNA amplification. Moreover, the possible occurrence of chimeric molecules in cells overproducing a specific protein, such as our study model, was predicted by the postulated amplification mechanism (1, 2, 18). The results of the present study constitute a confirmation of this prediction and validate the proposed mechanism. While we refer the reader to a previous study where the postulated model is described in detail (2), it can be briefly summarized as follows.

The amplification process starts with transcription of the antisense complement from a conventional mRNA template initiating at the 3’poly(A), possibly with the help of a uridylated protein (Fig. 1, Step 1). Generation of a complete antisense transcript requires the presence of an eligible RNA template and a compatible polymerase activity. The only major prerequisite for a potential RNA template appears to be the presence of the poly(A) segment at its 3’ terminus (1, 2, 19). The compatible polymerase activity is the RNA-dependent RNA polymerase, RdRp (20, 21). However, on its own, this apparently “omnipresent” core (2) activity has low processivity and terminates after generating only short transcripts (19). It appears that another activity, a “processivity co-factor” (2) that is induced in selective circumstances, is needed in addition to the core enzyme to efficiently produce the full-length transcripts. The resulting double stranded sense/antisense structure is then separated into singlestranded molecules by a helicase activity that mounts the poly(A) segment of the 3’poly(A)-containing strand (the sense-oriented strand) of the double helical structure and proceeds along this strand modifying, on average, every fifth nucleotide in the process (ref. 2; Fig. 1, Step 2). The 5’ poly(U)-containing antisense strand remains unmodified during and after the separation (2), this being essential for the production of a new sense strand since modifications could interfere with complementary interactions required in this process.

The vast majority of mammalian mRNA species contains 3’-terminal poly(A) segments. The notion that many, or possibly most, of them could be eligible templates for RdRp was suggested in our previous studies (1, 2). Subsequent observations by Kapranov et all showed a widespread synthesis of antisense RNA initiating, apparently indiscriminately, at the 3’ poly(A) of mRNA in human cells (19). This, apparently undiscerning, RdRp template eligibility of the bulk of mammalian mRNA species raises questions with regard to mechanisms underlying the manifestly stringent specificity of the mRNA amplification process as seen, for example, in erythropoietic differentiation (1, 2). The specificity of mRNA amplification appears to be determined at the 3’ terminus of an antisense transcript by its ability or inability to support production of a complementary sense strand molecule.

The generation of a sense strand on an antisense template occurs via the extension of the 3’ terminus of a self-primed antisense template and requires the presence within the antisense transcript of two spatially independent complementary elements (22). One of these is the strictly 3’-Terminal Complementary Element (TCE), the other is the Internal Complementary Element (ICE). These elements (Fig. 1, Step 3) must be complementary to a sufficient extent to form a priming structure but may contain mismatches and utilize unconventional G/U pairings. The generation of a sense strand also requires the thermodynamic feasibility, enhanced/enabled by the occurrence of two complementary and topologically compatible elements, of the antisense strand folding into a self-priming configuration.

Provided that a self-priming structure is formed, the 3’ end of the folded antisense strand is extended by RdRp into a sense-oriented molecule terminating with the poly(A) at the 3’end (Fig. 1, Step 4), thus generating a hairpin-structured chimeric intermediate consisting of covalently joined sense and antisense strands. The double stranded portion of the resulting structure is separated by a helicase activity invoked above, which mounts the 3’poly(A) of a newly synthesized sense strand component of the chimeric intermediate and proceeds along this strand in the 5’ direction modifying the molecule as it advances (Fig. 1, Step 5). When the helicase activity reaches a single stranded portion of the hairpin structure, it, or associated activities, cleave the molecule either within the TCE, at a TCE/ICE mismatch, or immediately upstream of the TCE; the cleavage occurs between the 5’ hydroxyl group and the 3’ phosphate (refs. 1, 2; red arrowhead, Fig. 1, Step 6).

Strand separation, in conjunction with the cleavage, produces two single-stranded molecules (Fig. 1, Step 7) one of which is a chimeric mRNA, the functional mRNA end product of amplification and the basis for defining this pathway as the “chimeric”. The chimeric nature of this end product is due to the presence at its 5’ end of a 3’-terminal segment of the antisense strand consisting, depending on the site of cleavage of the chimeric intermediate, of either the entire TCE or a portion thereof covalently attached, in a 5’ to 3’ orientation, to the 5’-truncated sense strand. This chimeric molecule is modified and 3’ polyadenylated; it cannot be further amplified because its antisense complement would be lacking the TCE but can be translated into either the conventional mRNA-encoded polypeptide (2) or its C-terminal fragment (18), depending on the extent of 5’truncation. The extent of 5’truncation of the sense strand component of chimeric end product is defined by the intramolecular position of the internal complementary element, ICE, within the antisense template and allows for conceptually distinct outcomes. Figure 1 illustrates the situation whereby the ICE of the antisense strand is located within its segment corresponding to the 5’ untranslated region, UTR, of a conventional progenitor mRNA. Consequently, the chimeric end product contains the entire protein coding region of a conventional mRNA and can be translated into the original, conventional mRNA-encoded, polypeptide (9). Additional translational outcomes reflecting alternative positions of the internal complementary element are discussed elsewhere (18).

The best, if not the only, identifier of the occurrence of mRNA amplification is a chimeric junction between sense and antisense components. Although the chimeric end product of amplification would be the most abundant source of the junction sequences, it may, for two reasons, not provide the best, or possibly even any, evidence for the occurrence of the amplification. First, as described earlier (2), the modified molecules could not be sequenced because modifications either impede the progression of reverse transcriptase, or interfere with complementary interactions, or generate inter- or intramolecular structures impassable by a polymerase. However, even if such a sequence could be obtained, it may not constitute good evidence. This is because the antisense component could be too short for a definitive identification or even indistinguishable from a sense-oriented sequence. Indeed, if there are no G/U pairings, if a strand-separating activity would cleave at the first mismatch within the TCE or if the TCE would be perfectly complementary to the ICE, the sequence of an antisense component of a chimeric RNA molecule would be identical to and indistinguishable from the sequence of the regular mRNA in the region of interest, and its only characteristic feature would be a 5’ truncation, insufficient for a definitive identification. The best “identifier”, therefore, appears to be a nucleotide sequence of a yet unmodified portion of uncleaved chimeric intermediate, containing sufficiently long and consequently unmistakably identifiable sense and antisense junction components. The chimeric fragments detected in the present study, where antisense components extend upstream from the junction site well beyond the loop portion, fit precisely this description.

Genes for all three chains of laminin 111 have multiple TSS positions (23, 24), most of which are inconsistent with the eventual generation of antisense molecules capable of self-priming within their segments corresponding to the 5’UTRs of mRNA because one of the complementary elements on the antisense strand is not 3’-terminal. Observations described above suggest a possibility that a shift in TSS positions can play an important regulatory role in defining the eligibility of a transcript for amplification, i.e. the ability of the antisense strand to fold into a self-priming configuration. The concepts of such a regulation are summarized in Figure 5.

**Figure 5.**
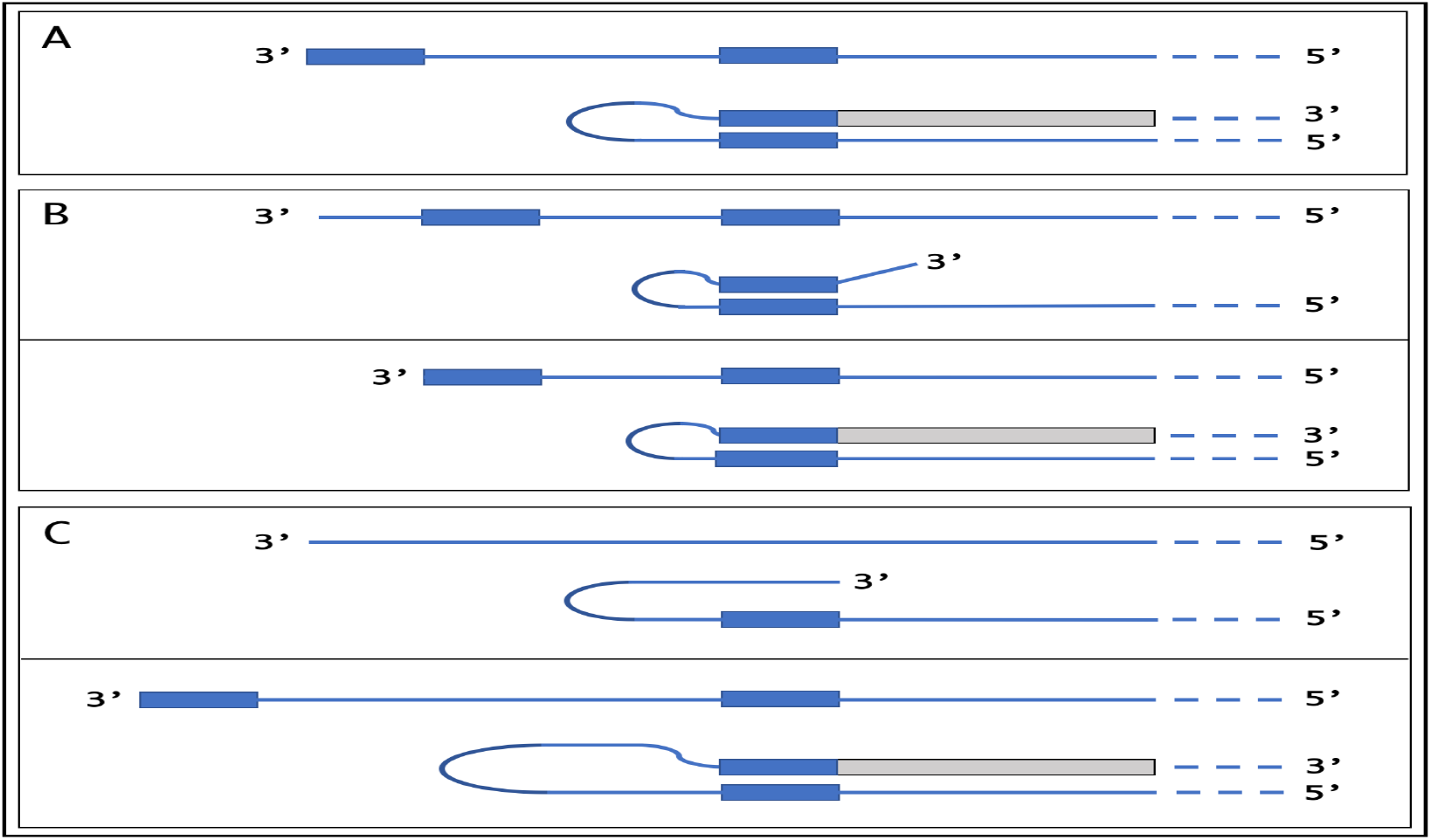
TSS shift as a potential regulator of the eligibility of an mRNA for the amplification process. Single line – 3’ terminus of the antisense strand. Filled grey boxes – sense strand. Filled blue boxes – topologically compatible complementary elements on the antisense strand. **A**: one of the complementary elements is 3’-terminal; folding results in a self-priming structure which is extended into the sense strand. **B**: both complementary elements are internal, no self-priming is possible; TSS shift in the downstream direction makes one of the elements 3’-terminal and allows self-priming and extension into the sense strand. **C**: there are no complementary elements, no self-priming can occur; TSS shift in the upstream direction generates complementary elements one of which is 3’-terminal and thus enables self-priming and extension. Note that processes depicted in panels B and C can occur in reverse resulting in a loss, rather than the acquisition, of the eligibility.

If the 3’-distant complementary element of the antisense strand is terminal (Figure 5, panel A), it can form a self-priming structure. If, however, both elements are internal (Figure 5, panel B), the downstream shift of the TSS position can make one of them a 3’-terminal and thus enable the self-priming. When, on the other hand, the 3’ segment of the antisense strand has no topologically compatible complementary sequences, an upstream shift of the TSS position (Figure 5, panel C) can generate such elements and make the transcript eligible for amplification. The events diagrammed in panels B and C of Figure 5 can also occur in reverse, making previously eligible mRNA ineligible for the amplification process.

The results of the present study indicate that mRNA encoding at least some ECM components may be amplified by the RNA-dependent mechanism and in a developmental stage-dependent manner. In this process, the occurrence of a mechanism regulating the eligibility of transcripts for amplification appears to be of a considerable importance. At the very early stage of the EHS tumor development, six days post-implantation, laminin 111 mRNAs encoding all three chains are expressed but are not amplified. This could be because the amplification machinery is not yet activated or because the transcripts are ineligible templates. At ten days after implantation, laminin is produced at high levels and its RNA transcripts, based on chimeric RNA detection, are amplified, i.e. the amplification machinery is active, whereas other major matrigel components, perlecan core protein, nidogen and collagen IV, are also expressed but at lower levels and there are no signs of amplification of their transcripts. The production of these basement membrane components picks up at the later stages of tumor development and there are good reasons to expect that their transcripts will also undergo amplification. It appears, therefore, that in a system with active amplification machinery the eligibility of transcripts for amplification is regulated. TSS shift can be such a regulatory mechanism for “TATA”-less genes which include both nidogen and collagen IV, as well as the majority of genes encoding ECM components.

Our previous results indicated that RdRp could transcribe the cap “G” of mRNA, despite its inverted orientation. Observations of the present study confirm this conclusion. The underlying arguments are based on the results with the a1 chain of laminin shown in the top panel of Figure 2 and are illustrated in Figure 6. If the cap “G” is not transcribed, the antisense strand terminates with the 3’ “u” (Fig.6, panel A; highlighted in blue) and its folding configuration would be as shown in Fig. 6, panel A. In this case, after the extension, the sense/antisense junction would consist of the “u/C” (highlighted in blue) as depicted in Figure 6, panel B. The experimental results, however, are different. They show that the sequence of the sense/antisense junction is, in fact, the “uc/C” (Fig. 6, panel C; highlighted in green). Since the genomic sequence upstream of the TSS cannot account for the additional 3’-terminal “c” (Fig.6, panel D; highlighted in green) in the antisense strand (23), the remaining possibility is that the “c” in question, unencoded in the genome, is a transcript of the cap “G” of the sense strand and that the antisense folding into a self-priming configuration occurs as shown in Figure 6, panel D.

**Figure 6.**
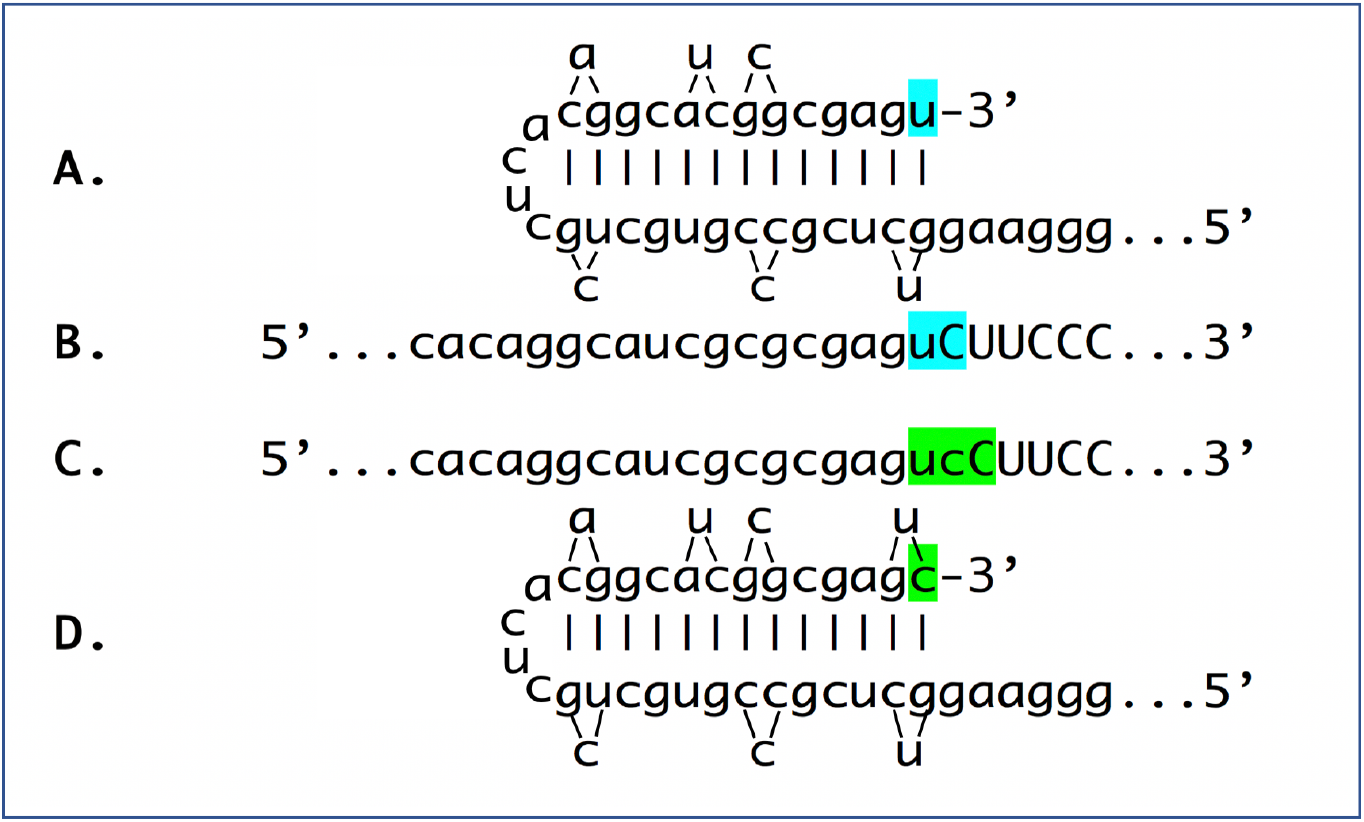
RNA-dependent RNA polymerase can transcribe the cap “G” of mRNA. Data shown is adopted from Figure 2, top panel. Uppercase letters – nucleotide sequence of the sense strand; lowercase letters – nucleotide sequence of the antisense strand. “u” highlighted in blue – 3’-terminal nucleotide of the antisense strand corresponding to the transcription start site of mRNA; “uC” highlighted in blue –the projected junction structure in the absence of the cap “G” transcription. “c” highlighted in green – transcript of the cap “G”; “ucC” highlighted in green – the resulting junction structure when cap “G” is transcribed. **A**: projected self-priming configuration of the antisense strand in the absence of the cap “G” transcription. **B**. Projected nucleotide sequence of the sense/antisense junction in the absence of the cap “G” transcription. **C**. Detected nucleotide sequence of the sense/antisense junction. **D**. Self-priming configuration of the antisense strand as defined by experimental results. Note that the genomic sequence upstream of the TSS cannot account for the additional 3’-terminal “C” in the antisense strand (23).

Until now, the occurrence of RNA-dependent amplification of mammalian mRNA was only shown for globin-encoding mRNA at rather extreme circumstances of terminal erythroid differentiation (1, 2). The finding of chimeric junction sequences for mRNAs encoding all three constituent chains of laminin 111 in a tissue producing extraordinarily large amounts of ECM proteins, indicates that such an amplification process may operate also during normal development when the production of large quantities of certain polypeptides is required. The determination of the extend of amplification of laminin mRNA in studied system must await the detection and quantification of the RNA end product of the amplification process but the possible range of amplification is suggested by the results obtained in the erythroid system. In the developing mammalian erythroid cell, the mass of conventional globin mRNA represents 0.01% of that of ribosomal RNA (10^4^ globin mRNA molecules, about 700 nucleotides each, and 10^7^ ribosomes, about 7000 nucleotides each, per cell; refs. 26, 27) whereas the mass of the apparent end product of globin mRNA amplification constitutes about 12% of that of ribosomal RNA (2), indicating a 1200-fold amplification. Considering the extreme nature of erythroid differentiation, this number probably represents the upper range of mammalian mRNA amplification.

It should be mentioned that the increase in number of molecules could be not the only consequence of mRNA amplification. The amplified mRNA appears to be heavily modified (2) and could behave in ways that are different spatially, qualitatively and quantitatively from those of conventional mRNA. For example, the amplification of mRNA species encoding secreted proteins, such as laminin, could overwhelm the ER, cause the ER stress and trigger cell death. It could be suggested, therefore, that nucleotide modifications of amplified mRNA, apparently acquired during strands separation (ref. 2; Fig. 1, step 5), may direct its translation and secretion of the resulting protein outside the ER, despite the presence of a secretion signal sequence. The initial ER stress resulting from increased transcription and subsequent translation of conventional mRNA encoding a secreted protein could be one of potentially multiple categories of the triggers of mRNA amplification processes. In this case, one can envision that a conventional overproduction of a secreted protein induces ER stress (known to result in the production and export of several transcription factors) which, in turn, activates mRNA amplification pathway thus facilitating overproduction and, in fact, relieving the ER stress since in the mRNA amplification process, a portion of conventionally produced mRNA molecules, used as templates for the production of antisense RNA, is apparently modified during strands separation (ref. 2; Fig. 1, step 2) and would be translationally processed outside of the ER, alongside the end product of mRNA amplification. Nucleotide modifications of amplified mRNA could also facilitate its cap-independent translation and even provide a quantitative advantage. In the erythroid system, the end product of globin mRNA amplification is not capped (it rather terminates with 5’OH group) yet it is efficiently translated (2).

In eukaryotes, RNA-dependent RNA polymerase was convincingly detected also in normal plant cells (28–30). Whereas these findings imply the possibility of RNA-dependent mRNA amplification, the occurrence of this process in plant organisms remains to be demonstrated.

As described elsewhere (18), the RNA-dependent mRNA amplification mechanism can produce not only mRNA molecules with the intact coding content of a conventional mRNA but may also generate 5’-truncated mRNA encoding C-terminal segments of conventional proteins, and, potentially, even unrelated polypeptides (11, 18). In this respect, the potential physiological significance of the mRNA amplification mechanism, with every conventional genome-originated mRNA molecule acting as a possible template, can hardly be overstated. Malfunctions of this process may be involved in pathologies associated either with the deficiency of a protein normally produced by this mechanism or with the overproduction of a protein or of its C-terminal fragment, as was proposed for the overproduction of beta amyloid in sporadic Alzheimer’s disease (11, 18).

## MATERIALS AND METHODS

EHS tumor tissue, removed from mice six or ten days post-implantation, was provided by MuriGenics. Inc. (CA). Tissue was reduced to a cell suspension by rubbing against and washing through a 70μm cell strainer. Pelleted cells were lysed in a buffer containing 30 mM Hepes, pH8.2; 50mM NacCl; 5 mM MgCL2; 1% NP40; 10% sucrose; 5 mM DTT and 1.5 units/μl RNase inhibitor (NEB). Nuclei were removed by centrifugation for 3 min at maximum speed in an Eppendorf centrifuge and the supernatant was vigorously mixed with 10 volumes of Trizol reagent. RNA was isolated from the Trizol mix as suggested by the manufacturer.

Note that RNA intended for studies of RNA amplification was extracted from an isolated cytoplasmic fraction, after removal of cell nuclei. This is because the bulk of amplified RNA, both end product and intermediates, is at least partially modified. It tends to bind to DNA (2) and would be removed if the isolation is attempted from total cell lysates. Since, in the absence of DNA, modified RNA binds to a predominant RNA species (2), cytoplasmic RNA was fragmented as described below prior to the rRNA depletion in preparation for the construction of sequencing libraries.

For rRNA depletion, total cytoplasmic RNA was fragmented by incubation for 12 minutes at 94°C in first strand synthesis buffer (NEB, kit E7770). Following rRNA depletion with NEBNext rRNA depletion kit ((E6310), RNA was purified with SPRI magnetic beads and used for preparation of sequencing libraries. Sequencing libraries were prepared using NEBNext Ultra II RNA library preparation kit for Illumina (NEB E7770) as suggested by the manufacturer with the following modifications: Step 1 (fragmentation and priming) was for 3 minutes; in step 2 (first strand synthesis) 8ul of the strand specificity reagent (NEB) was added instead of water and an additional incubation at 50°C for 15 minutes was included; after adaptor ligation and before PCR enrichment, an additional round of purification on SPRI beads was included; PCR enrichment was carried on for 7 cycles. After quality control, libraries were sequenced by the Harvard Biopolymers Facility using Illumina Next500 instrument. The initial analysis of the antisense folding patterns was carried out with the help of IDT oligonucleotides analyzing software. Sequencing data obtained were converted into blasting databases with the help of the Harvard Research Computing department and blasted against reference sequences at the stringency of word size 7. Sequences of interest were then extracted from the raw data and analyzed.

## ACKNOWLEDGEMENT

Authors are grateful to George Martin, Henry Lopez and MuriGenics Inc. for providing EHS tumor tissues, to New England Biolabs for gift of valuable reagents, and to Kathleen Keating and Harvard Research Computing for assisting with the data analysis.

## AUTHORS CONTRIBUTION

V.V. developed the underlying concepts, designed and performed experiments, analyzed data and wrote the manuscript. S.R. and B.O. contributed to conceptualization, discussions, data analysis and preparation of the manuscript. B.O. suggested the study model.

